# Transgenerational cues about local mate competition affect offspring sex ratios in the spider mite *Tetranychus urticae*

**DOI:** 10.1101/240127

**Authors:** Alison B. Duncan, Cassandra Marinosci, Céline Devaux, Sophie Lefèvre, Sara Magalhães, Joanne Griffin, Adeline Valente, Ophélie Ronce, Isabelle Olivieri

## Abstract

This preprint has been reviewed and recommended by Peer Community In Evolutionary Biology (https://doi.org/10.24072/pci.evolbiol.100051). In structured populations, competition for mates between closely related males, termed Local Mate Competition (LMC), is expected to select for female-biased offspring sex ratios. However, the cues underlying sex allocation decisions remain poorly studied. Here, we test for several cues in the spider mite *Tetranychus urticae*, a species that was previously found to adjust the sex ratio of its offspring in response to the number of females within the local population, i.e. a patch. We here investigate whether the offspring sex ratio of *T. urticae* females changes in response to 1) the current number of females in the same patch, 2) the number of females in the patches of their mothers and 3) their relatedness to their mate. Single females on patches produced similar sex ratios to those of groups of 15 females; their mothers had been in identical conditions of panmixia. The offspring sex ratios of females mated with their brother did not differ from those of females mated with an unrelated male. Females however produced a more female-biased offspring sex ratio if their mothers were alone on a patch compared to 15 other females. Thus, maternal environment is used as a cue for the sex allocation of daughters. We discuss the conditions under which the maternal environment may be a reliable predictor of LMC experienced by grand-sons.

## INTRODUCTION

In structured populations, dispersal of one sex can cause competition to be more intense between kin of the nonispersing sex than between kin of the dispersing one. This asymmetry in competition can cause a bias in the offspring sex ratio to reduce competitive interactions between brothers or sisters. If females are the dispersing sex, related males can compete for mates, a phenomenon termed Local Mate Competition (LMC). Female-biased offspring sex ratios are expected to evolve to reduce this competition between males, both through a decrease in the number of male competitors and an increase in the number of male interactions between brothers or sisters. competitors and an increase in the number of mates per son (Hamilton, 1967; Maynard Smith, 1978; Taylor, 1981; Charnov, 1982; Frank, 1985; Herre, 1985). In addition to LMC, for haplodiploid species, a more female-biased offspring sex ratio can arise because mothers that mate with a brother are more related to their daughters than they are to their sons (Greeff, 1996; Reece *et al*., 2004). Inbreeding does not change the relative relatedness of mothers to sons as they only receive genes from their mothers (Greeff, 1996).

Population structure can vary in time, and consequently so might the intensity of LMC and sib-mating. In such a situation, selection should favour females that facultatively (plastically) adjust their sex allocation in response to cues in the environment that reflect the intensity of LMC their sons will experience as well as whether they have mated with kin (Frank, 1985; Herre, 1985; Nagelkerke & Sabelis, 1996; Macke *et al*., 2011a). For example, females could adjust their offspring sex ratios in response to the number of females laying eggs (foundresses) in the same patch or to the level of relatedness of their mate. Consistent with expectations that females respond to current cues in the environment, more female biased offspring sex ratios are observed in patches formed by a single foundress as compared to patches founded by multiple foundresses, across a wide range of species (Shuker *et al*., 2005; West *et al*., 2005; Sato & Saito, 2007; Wang *et al*., 2015). Several studies, however, found that females do not adjust their offspring sex ratio in response to different numbers of foundresses on a patch (Innocent *et al*., 2007; Abe *et al*., 2014; Lievens *et al*., 2016). Moreover, there is currently no empirical support for the prediction that sib-mating results in more female biased offspring sex ratios than does random mating (Reece *et al*., 2004; Burton-Chellew *et al*., 2008).

Alternatively, LMC and inbreeding may change in a predictable fashion over generations. This scenario is predicted by haystack models of LMC in which sub-divided populations may persist over multiple generations, and relatedness between females sharing a patch may increase across generations (Nagelkerke & Sabelis, 1996). This increased relatedness can select for even more female biased offspring sex ratios due to more intense LMC (Frank, 1985; Gardner *et al*., 2009). If change in the sub-divided population structure is predictable through time, transfer of information from mothers to daughters could allow for a better adjustment of sex allocation.

Discrepancies among studies investigating sex allocation may be caused by the underlying cues being unknown and thus overlooked when designing experiments. In principle, these cues could stem from both the current or past environment, i.e. factors experienced while a female is laying her eggs, or those encountered as a juvenile, or even in previous generations. For example, females of the parasitoid wasp *Nasonia vitripennis* change their sex allocation in response to immediate cues in the environment, including the current number of foundresses, female odours and the presence of eggs in the environment (Shuker *et al*., 2005, 2007). In contrast, it is the environment that house wrens experience as juveniles, not the current environment, that affects the sex ratio of their own offspring: they produce a more female-biased offspring sex ratio when raised in a smaller brood (Bowers *et al*., 2017). Also, female mice exposed to stress *in utero* produce more daughters (Edwards *et al*., 2016).

Here we investigate the impact of immediate and past cues on sex allocation in the spider mite *Tetranychus urticae*. An earlier study showed that spider mites evolving under constant, high LMC lost the ability to facultatively adjust their offspring sex ratio, whilst those evolving under weaker LMC retained the ability for facultative sex allocation in response to the number of foundresses (Macke *et al*., 2011a). In that study, sex ratio was measured after mites had spent two generations on patches containing the same number of foundresses. Thus, both mothers and their daughters experienced cues indicating the same intensity of LMC for their sons. Therefore, the set-up did not allow disentangling the current vs past cue(s) used by females that retained the ability for facultative sex allocation. Other studies have shown that maternal cues impact a number of traits in spider mites including sex allocation, dispersal distance, fecundity and diapause (Oku *et al*., 2003; Magalhães *et al*., 2011; Bitume *et al*., 2014; Marinosci *et al*., 2015). *T. urticae* females also respond to current cues and produce more female-biased offspring sex ratios when there are fewer females sharing a patch (Wrensch & Young, 1978; Roeder, 1992) and they are able to recognise kin (Tien *et al*., 2011; Bitume *et al*., 2014).

We aim at uncovering the interplay between immediate and past cues in the environment in determining sex ratios in *T. urticae*. We investigate whether females alter their sex allocation in response to the number of females in the same patch (i.e. current indicators of LMC that their sons will experience) or the number of females in the patch of their mothers (i.e. past LMC conditions). Because sib-mating is negatively correlated with the number of foundresses on a patch, and a certainty in our high LMC treatments, we evaluate the effect of sib-mating on sex allocation in a separate experiment.

## MATERIAL AND METHODS

### Biological model

The two-spotted spider mite, *T. urticae* Koch (Acari: Tetranychidae), is a generalist hebivore (Migeon & Dorkeld, 2006-2017; Helle & Sabelis, 1985) with a short generation time (less than 15 days at 20°C on a suitable host; (Helle & Sabelis, 1985; Riahi *et al*., 2013). It has an arrhenotokous haplodiploid life cycle in which females produce haploid sons parthenogenetically and diploid daughters from fertilised eggs. After the egg stage, all individuals pass through a larval and two nymphal (protonymph and deutonymph) stages, with short quiescent stages between each, before reaching maturity. Mature females are approximately twice as big as, and rounder than, mature males. The sex ratios measured in this study are tertiary sex ratios defined as the proportion of adult males on a patch; offspring sex ratio is a trait of the mother.

### Study population

This study uses the same base population as Macke *et al*. (2011a). This population was collected from a cucumber greenhouse in Pijnacker, The Netherlands, in May 1994. Mites were transferred to a climate chamber at the University of Amsterdam where they were maintained on the same host plant. In September 2007, after ~240 months (~480 generations), approximately 5000 mites were sampled from the Amsterdam population to create two populations on cucumber plants at the University of Montpellier. Mite populations were maintained under the same conditions in a single climate chamber at 25°C ± 2°C, with a photoperiod of 14L:10D. A new, more genetically variable, base population was created in February 2011 by mixing and transferring approximately 200 adult females from the two previous populations. This new base population was maintained on 4 - 6 cucumber plants contained in a single plastic box (505 mm length × 310 mm width × 380 mm height) with a hole in the lid covered in a fine mesh to avoid condensation. Plants were watered every week and rotten plants removed and replaced with young, mite-free plants; plants were placed beside infested ones to facilitate mite dispersal.

To start the experiment (Generation 0), 12 independent samples from the base population consisting of 40 adult females each were created; this ensured that females seeding the experiment were of the same age. Each sample was placed on a whole cucumber leaf resting on water-saturated cotton wool in a plastic box (255 mm × 185 mm × 77 mm) to prevent both leaf dehydration and mite dispersal. Females were allowed to lay eggs and, 14 days later, their mature adult daughters were sampled to start the experiment ‘*sex allocation in response to LMC across generations’*. In the *‘sib mating’* experiment, slightly older daughters were used because the experiment was started 20 and 21 days after females started laying eggs in generation 0. In all experiments, mites were maintained in the same plastic boxes on leaves or leaf fragments placed on saturated cotton wool in climate chambers at 25 ± 2°C with a 16L:8D cycle. Prior to starting and during the experiments, we confirmed that our *T. urticae* populations were *Wolbachia-* free (see Supplementary Materials, Appendix 1 for details).

### Sex allocation in response to LMC across generations

This experiment investigated the sex allocation of females over two generations. In Generation 1 (the ‘maternal generation’), we measured the offspring sex ratios in response to high (1 foundress) versus low (15 foundresses) LMC experienced by laying females (these were the daughters of females from Generation 0). Each patch from Generation 0 contributed equal numbers of females, arbitrarily sampled, to the high and low LMC treatments. In Generation 2 (the ‘daughter generation’), we investigated the sex allocation under high LMC of daughters from females of Generation 1, which had experienced either high or low LMC. Thus, in Generation 2 we tested whether LMC experienced by mothers influenced the offspring sex ratios of their daughters (see Figure S1 for an outline of the experimental design).

#### Generation 1 (maternal generation)

In the low LMC treatment, a total of 360 females, in groups of 15 females, were placed on 60 cm^2^ cucumber leaf patches. We had 6 boxes, each containing 4 low LMC patches. In the high LMC treatment, 240 females were individually placed on a 4 cm^2^ leaf patch, and spread across 5 boxes, each containing 48 patches. Female density was the same for both low and high LMC conditions (0.25 females × cm^2^). Each box was divided into four quarters, each quarter containing 12 patches of single females in high LMC boxes and one patch of 15 females in low LMC boxes. This was done to get the same numbers of replicates (quarters) for each treatment with similar numbers of females laying eggs (12 for low LMC and 15 for high LMC). Note that we did not control for female mortality, i.e. there may have been fewer females contributing to the next generation in each treatment. In both LMC treatments, females laid eggs, and 14 days later we counted the number of emerging adults on each patch, and determined their sex using a stereo light microscope. Only measurements from a subset of 3 high LMC boxes and 2 low LMC boxes were included in the statistical analyses due to experimenter error.

#### Generation 2 (daughter generation)

Independently of LMC levels, 12 females were randomly collected from each quarter of each box from Generation 1 and individually placed under high LMC on a 4 cm^2^ leaf patch (same density as in Generation 1). Due to differential fecundity, some females from Generation 1 may have contributed more offspring to Generation 2 than did others. All patches were randomly spread across 6 boxes (48 patches per box). Note that some females in Generation 2 were sampled from patches that were not used in the analysis of sex ratio in Generation 1 due to experimenter error. Females were allowed to lay eggs and their offspring sex ratio was measured 14 days after transfer. In total, we measured the offspring sex ratio and number of emerging adults on 114 patches from 20 quarters in boxes testing the effect of low maternal LMC (5 - 6 females per quarter), and 116 patches, from 14 quarters, in boxes testing the effect of high maternal LMC (3 - 12 females per quarter). Some patches were lost due to female mortality before they laid eggs or to rotting of leaf patches.

### Sex allocation adjustment in response to sib mating

This experiment investigated whether females that mated with their brothers produced a more female biased offspring sex ratio than did females that mated with unrelated males. In total, 96 females sampled from Generation 0 were individually placed on 4 cm^2^ patches spread across 2 boxes (48 patches per box). This experiment was done in temporal blocks, over two days. Females were allowed to lay eggs and 9 days later quiescent daughters (2 from each patch, later allocated to each of the two treatments) were individually placed on 4 cm^2^ patches to ensure all mites assigned to our mating treatments were virgin. The same was done for emerged or quiescent sons. Mating pairs were formed three days later, when all quiescent females were adult. Females were assigned to one of two mating treatments: (1) the sib-mating treatment, in which one arbitrarily sampled brother was added to their patch or (2) the unrelated mating treatment, in which a male from another patch was added to the female’s patch. Males were removed after three days, females were allowed to lay eggs and 14 days later the sex ratio and number of emerging adults was measured. In total, we formed 24 patches with unrelated mate pairs and 28 patches with sibling mate pairs.

### Statistical analyses

For all analyses, except for the sex ratio of females from Generation 1 (see below), we analysed variation in the number of sons (offspring sex ratio) and the total number of offspring that emerged as adults using generalised linear mixed models, assuming a binomial distribution and a Poisson distribution, respectively. These models were performed in SAS using the GLIMMIX model procedure with the lsmeans statement to account for unbalanced designs (SAS script provided in Appendix 2, Supplementary Materials).

#### Sex allocation and offspring production in response to LMC across two generations

At Generation 1, the sex ratio was measured for single females for the high LMC treatment and for groups of 15 females for the low LMC treatment. To analyse variation in the number of sons across groups of females of similar sizes, we used a two-tailed permutation test, randomly sampling groups of 12 and 15 females (corresponding to the experimental design) under the null hypothesis of no effect of LMC. We generated 10,000 empirical values of the z statistics and could thus obtain the p-value for the effect of LMC on sex ratio. The observed statistic was generated using the number of sons produced in quarters of boxes by groups of 12 females for the high LMC treatment and by groups of 15 females for the low LMC treatment.

At Generation 1, we analysed variation in mean individual offspring production in each quarter as the number of offspring reaching adulthood in 14 days (this corresponds to ~4 days of laying eggs) accounting for LMC treatment as a fixed effect, and the box containing each quarter as a random effect nested within LMC treatment. At Generation 2, we tested the effect of the LMC conditions experienced by the mother at Generation 1 on male production and offspring production. We accounted for the LMC treatment of the mothers as a fixed effect, and quarter from which each female originated as a random factor nested within LMC treatment.

#### Sex allocation adjustment in response to sib mating

In the sib-mating experiment, we accounted for the mating treatments of the mothers as a fixed effect, and the patch of each mother and temporal block as random factors. Note that block could not be included in the analysis for offspring production because the model did not converge.

## RESULTS

In all experiments, mean (± 1 SE) offspring per female was not affected by the intensity of LMC in current (19.01 ± 1.630 for high LMC; 20.16 ± 0.920 for low LMC; F _1, 3_ = 0.13, p = 0.7397) or past generations (13.05 ± 0.860 for high LMC; 14.79 ± 0.744 for low LMC; F _1, 32_ = 2.37, p = 0.1335), or by the relatedness between mates (14.54 ± 1.088 for sib-matings; 16.42 ± 0.934 for unrelated matings; F 1, 21 = 2.95, p = 0.1005). The absence of an effect on offspring production ensures that the estimates of sex-ratio have the same precision between LMC and sib-mating treatments.

### Sex allocation in response to LMC across two generations

At Generation 1, females did not modify their offspring sex ratio according to the number of other females in the same patch (p = 0.938; Figure 1A). In contrast, at Generation 2, the maternal environment affected the sex allocation of their daughters, as females produced a less female biased offspring sex ratio if their mothers had experienced low, compared to high, LMC (F _1, 32_ = 10.98, p = 0.002, Figure 1B).

**Figure 1.**
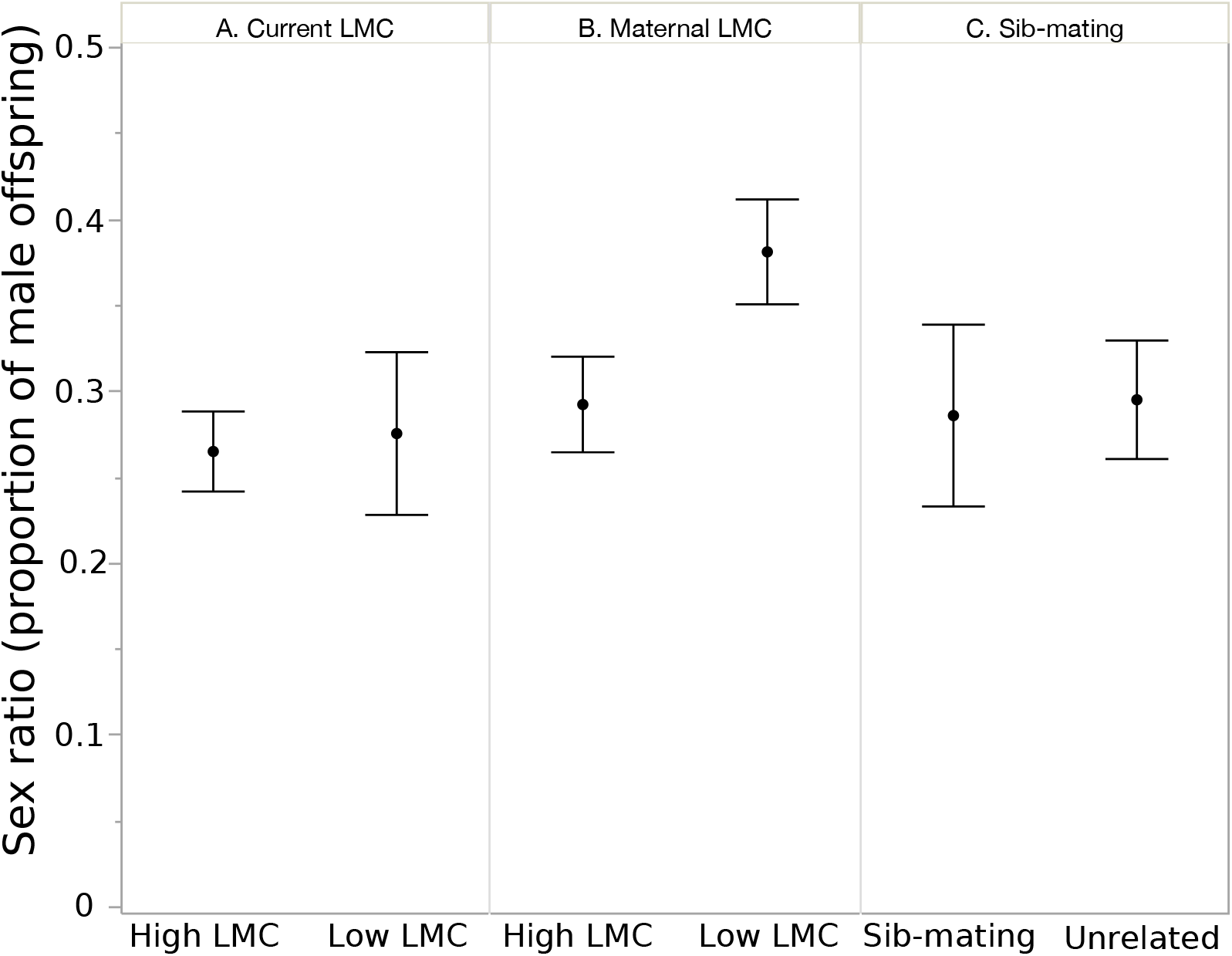
Proportion of male offspring for (A) females experiencing high (1 female) or low (15 females) LMC, (B) females whose mothers experienced high or low LMC and (C) females mated with their brother or an unrelated male. Figures show empirical means ± 2 standard errors (confidence interval).

### Sex allocation adjustment in response to sib-mating

Mating with a brother or an unrelated male did not change sex allocation (F _1, 21_ = 0.06, p = 0.82; Fig. 1C). This means the effect of high LMC over two generations on sex allocation are not due to sib-mating.

## DISCUSSION

We found that sex allocation in the spider mite *T. urticae* does not depend on the number of females on the same patch or on the relatedness between mates. Instead, we found that females alter their offspring sex ratio according to the number of females on their mother’s patch, showing that the maternal environment may be important for sex allocation. Additionally, none of the treatments-current or maternal levels of LMC, or mating with a sibling-impacted the number of offspring that were produced over a ~ 4 day period and reached adulthood.

Female *T. urticae* did not produce a more female-biased sex ratio when they mated with their brother compared to an unrelated male. As spider mites do recognise kin (Tien *et al*., 2011; Bitume *et al*., 2013), this calls for an explanation. One possibility may be that there is sexual conflict over the optimal offspring sex ratio (Shuker *et al*., 2009; Macke *et al*., 2014). In haplodiploid species, males only pass their genes to the next generation via daughters, thus it is always beneficial for males to produce a more female-biased offspring sex ratio, whereas for females the optimal offspring sex ratio depends on levels of LMC (Shuker *et al*., 2009; Macke *et al*., 2014; Shuker & Cook, 2014). Since *T. urticae* males can manipulate females to produce more female-biased offspring sex ratios (Macke *et al*., 2014) females may not alter their sex allocation in response to any cue provided by their mate as it may not be selectively advantageous.

As found in a number of other species (Reece *et al*., 2005; Innocent *et al*., 2007; Abe *et al*., 2014) we did not find that female *T. urticae* alter their sex allocation in response to the number of other females in the same patch (but see Wrensch & Young, 1978; Roeder, 1992). For example, in *Nasonia vitripennis* other cues, specifically egg presence and the relative number of eggs a female contributes to a patch, had a stronger influence on sex allocation than did the number of foundresses (Shuker & West, 2004; Burton-Chellew *et al*., 2008). It could be, for our population under our experimental conditions, that the number of females in the current patch does not provide reliable information about the LMC level that the offspring will experience. This may be because (a) females often move away from the patch or (b) females have high mortality during the egg laying period. In spider mites, the second hypothesis is possible if females are on low-quality plants (Magalhaes *et al*., 2007; Marinosci *et al*., 2015).

It may be that females respond to density and not the absolute number of females. In studies that found an effect of foundress number on LMC, density was not controlled (Wrensch & Young, 1978; Roeder, 1992), but we did control for density in our experiment. Alternatively, the absolute number of females on a patch could have been too low to elicit a response. Spider mites use mechanical and chemosensory processes via web and faeces to detect conspecifics (Royalty *et al*., 1992; Clotuche *et al*., 2014). *T. urticae* can also use volatiles released by infested plants to detect conspecifics, but at densities much higher than those in our experiment, and on whole plants, and not leaves as here (Pallini *et al*., 1997). These cues may not have been available in our experiment with low densities of females.

In Generation 2 densities were also identical to those of Generation 1 and between treatments, and here we did detect a difference in sex allocation. Adult females whose mothers experienced low LMC produced less female biased offspring sex ratios. In both Macke *et al*. (2011a) and our experiment sex allocation is more female biased when females experienced at least two generations of LMC. In both experiments sex allocation could be in response to the number of foundresses in the mother’s patch, or the number, or relatedness, of individuals in the juvenile environment.

Our finding could be a true maternal effect in sex allocation, meaning that the mother’s experience of her environment influences her offspring’s phenotype (Mousseau & Fox, 1998). Maternal effects have a role in determining many offspring traits including offspring size and juvenile survival (Pick *et al*., 2016), resistance to parasites (Little *et al*., 2003) and competitive ability (Bentz *et al*., 2016). In *T. urticae* maternal effects partly determine offspring phenotypes for dispersal distance (Bitume *et al*., 2014), diapause incidence (Oku *et al*., 2003) and offspring life-history traits (Magalhães *et al*., 2011; Marinosci *et al*., 2015). To the best of our knowledge, parental effects have not been previously shown to play a role in determining offspring sex allocation. An exception may be studies in which stress during the embryonic period influences offspring sex ratio. For example, female mice produce a more female biased offspring sex ratio if they experience stress *in utero* (Edwards *et al*., 2016), and paternal incubation temperature influences offspring sex ratio in a lizard (Warner *et al*., 2013). Whether these can be considered as maternal (or paternal) effects is debatable, as the developing individual is itself experiencing the stress. If our results are due to a maternal effect the same one might be responsible for the results observed in the selection experiment by Macke *et al*. (2011a). Indeed, in that experiment, mothers and daughters experienced the same predictable LMC level, so sex ratio adjustment may be explained by the maternal environment.

If female sex allocation in *T. urticae* is in part influenced by maternal cues for LMC this could come about via the manipulation of a daughter’s egg production. Egg size is thought to be one way by which female *T. urticae* could plastically adjust their sex allocation, as fertilisation probability increases with egg size, bigger eggs being more likely to develop into females (Macke *et al*., 2011b). Furthermore, *T. urticae* evolving under high LMC produce larger eggs and a more female biased sex ratio (Macke *et al*., 2012). Female *T. urticae* possess a single ovary within which eggs develop and are fertilised (Feiertag-Koppen & Pijnacker, 1982). Female larvae that have recently emerged from their eggs possess cells in their ovaries undergoing meiosis (Feiertag-Koppen & Pijnacker, 1982; Mothes-Wagner, 1984); the conditions or resources of the mothers could influence the development and size of their daughters’ eggs, which could translate into a given sex ratio.

Another possibility is that the juvenile environment acts as a possible proximal cause of sex allocation, which itself depends on maternal LMC. When mothers experience high LMC, and are alone on a patch, females will develop with their siblings whereas under low LMC, juveniles will share their patch with both related and unrelated individuals. Furthermore, although we controlled for density across treatments, under low LMC there is a higher absolute number of juveniles than there is under high LMC. Offspring sex ratio has been linked to the environment a mother experiences during the juvenile period in wrens and voles; when mothers experienced juvenile resource limitation they produced less male biased offspring sex ratios (Helle *et al*., 2012; Bowers *et al*., 2017). Nevertheless, if females respond to cues experienced whilst juvenile and before they disperse to find their own patch, the conclusion remains that factors due to maternal LMC influence a daughter’s sex allocation.

Why, then should individuals use maternal cues, or cues available from the juvenile environment, rather than those present in the immediate environment to adjust the sex ratio of their offspring? It might be that our observed response is not adaptive. Indeed, maternal effects do not necessarily increase offspring fitness (Sheriff & Love, 2013; Barks & Laird, 2015). However, it might be adaptive if levels of LMC change in a predictable fashion across generations (for example, increasing or decreasing LMC) allowing a female to integrate information provided by mothers (Mousseau & Fox, 1998). It has been hypothesised that *T. urticae* life-history follows predictable population groupings across multiple generations on the same host plant (Nagelkerke & Sabelis, 1996). If this is the case, maternal cues may provide reliable information about the current state of a population, especially if relevant cues are not available in the patch. Mitchell (1972) showed that patches in experimental populations can be relatively small (1 cm^2^) compared to the size of a leaf, and the patch sizes we used. Patch size and population structure remain to be measured in natural populations.

We can conclude from our experiment and others that multiple cues act together to influence sex allocation decisions, with the contribution of each depending on context. Maternal cues (maternal LMC or juvenile environment) may provide a more integrative measure of population structure, which females use for their sex allocation decisions at low densities. Clearly, more studies are needed to assess the relative value of cues informing on population structure and on the precise scale at which competition for mates actually occurs.

## ACKNOWLEDGEMENTS

We would like to thank David Carbonell for growing and maintaining plants for this experiment. This project was funded by a joint grant from the Agence Nationale de la Recherche and the Fundação para a Ciência e a Tecnologia to IO and SM (FCT-ANR/BIA-EVF/0013/2012). This is ISEM contribution number ISEM 2018-096.

## DATA AVAILABILITY

Raw data are available as supplementary files.

